# Into the tropics: phylogenomics and evolutionary dynamics of a contrarian clade of ants

**DOI:** 10.1101/039966

**Authors:** Michael G. Branstetter, John T. Longino, Joaquín Reyes-López, Ted R. Schultz, Seán G. Brady

**Affiliations:** Department of Biology, University of Utah, Salt Lake City, UT 84112, USA; Department of Entomology, National Museum of Natural History, Smithsonian Institution, Washington, D.C., 20560, USA; Departamento de Botánica, Ecología y Fisiología Vegetal, Universidad de Córdoba, Córdoba, Spain

**Author notes:** Corresponding Author: Michael G. Branstetter, Department of Biology, University of Utah, 257 S 1400 E, Salt Lake City, UT 84112, USA.

**Keywords:** ants, biogeography, diversification, latitudinal diversity gradient, macroevolution, out-of-the tropics, ultraconserved elements

## Abstract

**Aim:** The standard latitudinal diversity gradient (LDG), in which species richness decreases from equator to pole, is a pervasive pattern observed in most groups of organisms. Despite its commonness, an increasing number of non-conforming lineages have been identified, presenting a challenge for general explanations of the standard LDG. Although problematic, documenting and studying these contrarian groups can help us to better understand LDGs generally. To that end we identify the ant tribe Stenammini, a diverse lineage with over 400 species, as a contrarian clade, and we attempt to explain the group’s atypical diversity pattern using a historical approach. We evaluate the following alternative hypotheses: time-for-speciation/center-of-origin (TFS/COO), niche conservatism, and differences in diversification rate.

**Location:** Global.

**Methods:** We examine the shape of the LDG in Stenammini by plotting latitudinal midpoints for all extant species. We then infer a robust phylogeny using a phylogenomic approach that employs ultraconserved element loci and we use the phylogeny to estimate divergence dates, biogeographic history, and ancestral habitats. We also test for diversification rate heterogeneity across the tree and among lineages within the tribe.

**Results:** Stenammini has a skewed inverse latitudinal diversity gradient with an extratropical richness peak in the northern temperate zone. Our phylogenomic approach resulted in a robust phylogeny revealing five major clades and several instances of non-monophyly among genera (*Goniomma*, *Aphaenogaster*). The tribe and most major lineages originated in the temperate zone and inhabited temperate niches. Crown Stenammini dates to 52 Ma (Eocene Climatic Optimum) and most major lineages appeared soon after during a period of global cooling. Despite its temperate origin, the group invaded the tropics at least six times, but failed to diversify greatly there. Across the tree diversification increased from 17.2-1.9 Ma following the Mid-Miocene Climatic Optimum, and among lineages there was a rate increase in Holarctic *Aphaenogaster* + *Messor* just prior to 17.2 Ma.

**Main Conclusions:** Our results suggest that time, niche conservatism, and increased diversification have all contributed to the inverse latitudinal gradient in Stenammini. Among these processes, niche conservatism may be less important given that the tribe has dispersed to the tropics multiple times.

## INTRODUCTION

The observation that species richness decreases from equator to pole is the oldest and best-known pattern in biogeography (Hawkins, 2001; Willig et al., 2003). Although not a consistent pattern throughout the history of life on Earth, the current latitudinal diversity gradient (LDG) has existed for at least the last 30 million years and is pervasive (Mannion et al., 2013). The standard LDG occurs in most fauna and flora, in both terrestrial and marine realms, and across a range of spatial and temporal scales (Willig et al., 2003; Hillebrand, 2004; Field et al., 2009; Brown, 2014). Hypotheses attempting to explain the standard LDG abound, yet a broad consensus as to which hypothesis, or set of hypotheses, best predicts the gradient remains a topic of active debate (Pianka, 1966; Rohde, 1992; Willig et al., 2003; Wiens & Donoghue, 2004; Jablonski et al., 2006; Brown, 2014).

A challenge to understanding the standard LDG has been the discovery of a small but growing number of lineages that exhibit inverse diversity gradients, in which species richness peaks in extratropical regions and decreases toward the equator. These contrarian clades occur in a variety of marine and terrestrial groups, including marine bivalves (Krug et al., 2007), pelagic seabirds (Chown et al., 1998), ichneumonid wasps (Owen & Owen, 1974), and lampropeltinine snakes (Pyron & Burbrink, 2009). Seemingly problematic, discovery of these groups has actually improved our understanding of diversity gradients, because it has made it necessary to explain these non-standard cases using the same fundamental mechanisms (i.e., speciation, extinction, and dispersal) that underlie the standard LDG. Thus, a productive goal in biogeography should be to identify contrarian clades and to understand the ecological and evolutionary processes that have caused them to be different. Here, we demonstrate that a major group of ants is a contrarian clade and we infer a robust phylogeny of the group to test several hypotheses that might account for the group’s atypical diversity pattern.

Ants (Hymenoptera, Formicidae) are the most successful group of eusocial insects on Earth. They occur in most terrestrial ecosystems and are often ecologically and numerically dominant (Hölldobler & Wilson, 1990). As a whole, ants exhibit the standard LDG (Kaspari et al., 2003), with the Neotropics being particularly important in the group’s evolutionary history (Moreau & Bell, 2013). Several ant lineages, however, are predominately temperate in distribution and exhibit obvious adaptations to living in colder, seasonal environments. One of the largest of these groups is the myrmicine ant tribe Stenammini (sensu Ward et al., 2015)

Stenammini is a monophyletic tribe that currently includes seven genera and over 400 extant species (Bolton, 2015). It occurs across the globe, being absent from only Amazonia, temperate South America, and permanently frozen regions. Considering type localities of described species, the majority of the tribe’s diversity occurs in the temperate zone of the northern hemisphere, with less than one quarter of the species known from the New and Old World tropics, a pattern that strongly suggests an inverse latitudinal gradient. Ecologically, the tribe exhibits a variety of life history strategies, but the group is best known for several genera that have radiated in arid landscapes as seed harvesters. These lineages are the genera *Messor, Veromessor, Novomessor, Goniomma*, and *Oxyopomyrmex*. The remaining two genera in the tribe, *Stenamma* and *Aphaenogaster*, are less specialized in their diets and are more strongly associated with mesic forest habitats.

Among the seed harvester lineages the genus *Messor* is the most diverse. It includes over 100 species and is distributed across the Palearctic and in arid regions throughout Africa. The genus *Veromessor* includes only nine species and is restricted to arid regions of the western United States and west-central Mexico (Johnson, 2001). This genus was previously synonymized with *Messor* based on its deceptively similar seed-harvester morphology (Bolton, 1982), but recent molecular analyses revealed that the similarities are convergent and that the North American species are not closely related to *Messor* (Brady et al., 2006; Ward et al., 2015). The genus *Novomessor* includes only three species and occurs in the southwestern USA and northern Mexico. It was previously synonymized with the genus *Aphaenogaster* (Bolton, 1982), but was recently resurrected based on phylogenetic analyses that included molecular data (Demarco & Cognato, 2015). Because it is less specialized and scavenges on a diverse range of food items *Novomessor* does not have the seed-harvester gestalt, but it is adapted to arid environments and forages on seeds as a significant part of its diet (Whitford et al., 1980; Johnson, 2001). The last two seed-harvester genera are *Goniomma* and *Oxyopomyrmex*. Both of these have large eyes and forage for seeds at night. Together they include about 20 species and are restricted to arid habitats around the Mediterranean region (Salata & Borowiec, 2015).

Of the remaining two members of Stenammini, *Aphaenogaster* is the largest and most taxonomically problematic genus in the tribe. It currently includes over 180 described species, distributed mostly throughout the Holarctic, but with some species diversity in the Neotropical (Mexico, Central America, Haiti), Afrotropical (Madagascar), Indomalayan, and Australasian regions. Within Australia, a few species penetrate into the southern temperate zone. Although the genus *Novomessor* was recently resurrected and removed from *Aphaenogaster*, considerable uncertainty remains as to whether *Aphaenogaster* is monophyletic (Ward et al., 2015). The group shows considerable morphological diversity and includes a number of genus-level names that are currently synonymized under *Aphaenogaster* (Bolton, 2015). Ecologically, the majority of species are scavengers that inhabit forested regions in the temperate zone, but a number of species have penetrated into arid and rainforest habitats in both the New and Old Worlds. Morphologically, one of the more striking features in the genus is the tendency for tropical species to become large and elongate, with the posterior end of the head extending into a long neck (e.g., *A. dromedaria*).

The genus *Stenamma* is the third largest member of the tribe with just over 80 described species. It occurs throughout the Holarctic and partway into the Neotropics, reaching as far south as northern South America (Branstetter, 2009, 2012, 2013). The genus differs from the rest of the tribe in that it is composed primarily of small-sized, cryptic ants that are most often found in the leaf litter of forested regions. Recent molecular work has demonstrated that the genus is monophyletic and that it is divided into equally diverse Holarctic and Neotropical clades (Branstetter, 2012). It also determined that the genus likely originated in the Nearctic and dispersed into the Neotropics, where it radiated in montane wet forests. Although the genus does not have an inverse latitudinal gradient, it does represent a contrasting example to the out-of-the tropics hypothesis, which proposes that most lineages evolved in the tropics and dispersed out (Jablonski et al., 2006).

The fossil record for Stenammini is seemingly rich (Bolton, 2015), including 18 putative species of *Aphaenogaster*, one species of *Stenamma*, one species of *Messor* (currently an unresolved junior homonym), and four species of a possibly related genus *Paraphaenogaster*. However, this richness belies the fact that the placement of many of these fossil taxa within the tribe, or within genera in the tribe, is dubious, largely due to uncertainty in both diagnostic characters and generic boundaries. Considering reliable fossils, the tribe has been documented from the Eocene Period onward, with records from various Eurasian deposits, including Baltic Amber (Wheeler, 1915), and several New World deposits, including the Florrisant Shale of Colorado (Carpenter, 1930), and the Dominican and Chiapas Ambers (De Andrade, 1995). The overall record suggests the tribe has been present throughout the Holarctic since the Eocene and in Central America since the late Oligocene.

In the present study we demonstrate that the ant tribe Stenammini is a biogeographic contrarian, exhibiting an inverse latitudinal gradient. Following this observation we take a historical approach, employing phylogenomic techniques and a variety of tree-based comparative methods (e.g. divergence dating, ancestral area and habitat reconstruction, and diversification rate-shift tests) to assess alternative hypotheses that might explain the tribe’s atypical diversity pattern. Specifically, we consider three hypotheses that have been used to explain LDGs and that can be evaluated using tree-based approaches. These are the time-for-speciation/center-of-origin (TFS/COO) hypothesis (Schluter, 1993; Wiens & Donoghue, 2004; Stevens, 2006), the niche conservatism hypothesis (Wiens & Donoghue, 2004; Wiens & Graham, 2005), and the diversification rate hypothesis (Rohde, 1992; Cardillo et al., 2005; Weir & Schluter, 2007). The TFS/COO proposes that species richness is primarily a function of the time a particular lineage has been present in a region. Thus, if a lineage has more species in the temperate zone, the prediction would be that the lineage originated in the temperate zone and has had more time to evolve there. The niche conservatism hypothesis proposes that niche evolution is rare, and a lineage with more temperate diversity should have a temperate ancestor and few transitions into the tropics and into tropical habitats. Lastly, the diversification rate hypothesis proposes that differences in richness are due to differences in speciation and extinction rates. Under this hypothesis, the region with more species should have either a higher speciation rate or a lower extinction rate.

## MATERIALS AND METHODS

### Assessing the Diversity Gradient

To examine the latitudinal diversity gradient in Stenammini we estimated the latitudinal midpoint for all extant described species in the tribe (see Appendix S1). This was carried out by inspecting the distribution of every species using an online database of ant species distributions (www.antmaps.org; Guénard et al., 2015) and recording the northernmost and southernmost point for each. In cases where only an administrative district was reported (country, state, *etc.*) rather than a point record, we estimated the latitude as either the midpoint of the country, or, alternatively, of a biogeographically sensible location. Using the midpoint latitudes we classified species as having either tropical or extratropical distributions and we generated a histogram of all points in R v3.2.2 (R Core Team, 2015).

### Phylogeny Taxon Selection

We selected eight outgroup and 92 ingroup species for inclusion in our study (see Appendix S2 for voucher information). Within Stenammini we included all genera and a selection of exemplar species from all regions where each genus occurs. For *Stenamma*, we included 23 species from the Neotropics and 14 species from the Holarctic, including a rare species from Morocco (*S. punctiventre*) that has not been sampled previously. We sampled all three species of the genus *Novomessor* and eight out of the nine species of *Veromessor*. We sampled five out of eight species of *Goniomma* and three out of 12 species of *Oxyopomyrmex*. For *Messor*, we only sampled six species, but these included taxa from southern Africa, Europe, and Asia. For *Aphaenogaster* we sampled 12 tropical species, nine Nearctic species, and nine Palearctic species. The tropical species include three from the Neotropics, five from Madagascar, one from Borneo, two from New Guinea, and one from Australia. One interesting species of uncertain affinity that we were unable to sample is *Aphaenogaster relicta*. Recorded from Haiti, it is the only known stenammine species to occur in the Caribbean. Despite several recent collecting trips to Hispaniola by colleagues, fresh material of this species was not acquired.

### Phylogeny Data Generation

To generate a robust phylogeny we employed a recently developed phylogenomic method that combines target enrichment of ultraconserved elements (UCEs) with multiplexed next-generation sequencing (Faircloth et al., 2012, 2015). The approach is preferable to more traditional Sanger sequencing because it results in orders of magnitude more data in less time and at less cost. The steps for conducting the lab work are the following: DNA extraction, library preparation, UCE enrichment, and sequencing. The protocols are outlined in Faircloth et al. (2015) and given in detail below. We extracted sequence data for 12 species from Faircloth et al. (2015) and generated new sequence data for the remaining taxa.

For all newly sampled taxa, we extracted DNA using Qiagen DNeasy Blood and Tissue kits (Qiagen Inc., Valencia, CA). We measured DNA concentration using a Qubit 2.0 fluorometer (Life Technologies) and we input up to 500 ng of DNA into shearing and library preparation. We sheared DNA to an average fragment distribution of 400-600 bp (verified on an agarose gel) using a Qsonica Q800R sonicator (Qsonica LLC, Newton, CT).

Following sonication, we constructed sequencing libraries using the standard Library Preparation Kit or the newer Hyper Prep Kit from Kapa (Kapa Biosystems, Wilmington, MA). We performed all reactions at ¼ volume except for the PCR step, which we assembled at full volume (50 μL). During library preparation we added custom, single-indexing Truseq-style barcode adapters to most samples (Faircloth & Glenn, 2012). However, for a few libraries, we used a newer, dual-indexing set of adapter-primers called iTru (Faircloth & Glen, pers. comm.). These adapters are analogous to the Illumina TruSeq HT system and are added to the libraries during the PCR step of library preparation. We assessed success of library preparation by measuring DNA concentration with the Qubit flourometer and by visualizing the libraries on an agarose gel. For a subset of the samples, we removed adapter-dimers by performing a 0.7-0.8X SPRI-bead cleaning.

For UCE enrichment we made pools at equimolar concentrations containing 6-10 libraries. We adjusted pool concentration to 147 ng/μl using a vacuum centrifuge and we used a total of 500 ng of DNA (3.4 μl of each pool) in each enrichment. We performed enrichments using a custom RNA bait library developed for Hymenoptera (Faircloth et al., 2015) and synthesized by MYcroarray (MYcroarray, Ann Arbor, MI). The probe set includes 2,749 probes targeting 1,510 UCE loci. We enriched each pool of samples following the manufacturer’s protocol, except that for some pools, we left streptavidin beads bound to enriched DNA during PCR (on-bead PCR). The enrichment was performed at 65°C for a period of 24 hours.

We verified enrichment success with qPCR by comparing amplification profiles of unenriched to enriched pools for several UCE loci. After verification, we used qPCR to measure the DNA concentration of each pool and we then combined all pools into an equimolar final pool. At most we combined 110 samples together into the same sequencing lane. To remove fragments that were either too large or too small for sequencing, we size-selected pools from 300-800 bp using the Blue Pippin size selection instrument (Sage Science, Beverly, MA). We mailed size-selected pools to either the UCLA Neuroscience Genomics Core, Cornell’s Institute of Biotechnology, or the High Throughput Genomics Facility at the University of Utah, and we had the samples sequenced as single lanes on an Illumina HiSeq 2500 (2x150 Rapid Run, or 2x125 v4 Run).

### Bioinformatics & DNA Matrix Preparation

After sequencing, technicians at the sequencing centers demultiplexed and converted the data from .bcl to .fastq format. Starting with the .fastq files, we performed all initial bioinformatics steps, including read cleaning, assembly, and alignment, using the software package PHYLUCE version 1.5 (Faircloth et al., 2012; Faircloth, 2015). Raw reads were cleaned and trimmed using *illumiprocessor* and cleaned reads were assembled *de novo* using the script *phyluce_assembly_assemblo_trinity*, which is a wrapper around the assembly program Trinity v2013-02-25 (Grabherr et al., 2011). We selected Trinity for read assembly because Faircloth et al. (2015) demonstrated that it creates better assemblies with hymenopteran UCE data than other popular assembly programs. After assembly, we mapped Trinity contigs to UCE loci using *phyluce_assembly_match_contigs_to_probes*, with the *min-coverage* and *min-identity* options set to 50 and 70, respectively. It should be noted that these less stringent settings increased the average number of captured UCE loci by several hundred loci as compared to the default settings in *phyluce*. Next, we used *phyluce_assembly_get_match_counts* and *phyluce_assembly_get_fastas_from_match_counts* to create a monolithic fasta file containing all taxa and sequence data. During these steps we used the *incomplete-matrix* flag to allow matrices to have missing data. We aligned all loci individually using *phyluce_align_seqcap_align*, which runs the alignment program MAFFT (Katoh et al., 2002). We used the *no-trim* and *incomplete-matrix* options during alignment to prevent the data from being trimmed and to include loci with missing data. We also adjusted the *min-length* and *proportion* options to 20 and 0.50, respectively. After alignment, we used *phyluce_align_get_gblocks_trimmed_alignments_from_untrimmed* to run the trimming program GBLOCKS (Castresana, 2000; Talavera & Castresana, 2007) on all aligned loci. We reduced the stringency of the default settings in GBLOCKS by adjusting the b1, b2, b3, and b4 options to 0.5, 0.5, 12, and 7, respectively. These settings were manually optimized by trial and error and visual inspection of alignments.

### Phylogenetic Inference

To reconstruct a robust phylogeny of the Stenammini we employed maximum likelihood (ML), Bayesian (BI), and species tree (ST) methods. We started by investigating the effect of missing data. To do this, we created several sets of filtered alignments using the PHYLUCE script *phyluce_align_get_only_loci_with_min_taxa*. This script removes loci that have fewer than a set percentange of the total number of taxa. We filtered the alignments for 100, 99, 95, 90, 75, 50, and 25% taxa present and we generated alignment stats for each locus set using the *phyluce_align_get_align_summary_data* and *phyluce_align_get_informative_sites* scripts. We then generated concatenated matrices for each locus set using the *phyluce_align_format_nexus_files_for_raxml* script. With the exception of the 100 and 99% filtered matrices (these included very few loci), we analyzed all concatenated matrices with RAxML v8.0.3 (Stamatakis, 2014), performing a rapid bootstrap analysis (100 replicates) plus best tree search (option “–f a”) using GTR+Γ as the model of sequence evolution. The “best” filtering level was determined subjectively by assessing matrix completeness, topology, and bootstrap scores. The best filtering level was selected at 90% and this resulting set of alignments was used for all subsequent analyses.

After filtering loci for missing data we carried out ML and BI analyses on the concatenated matrix using the programs RAxML v8.0.3 (Stamatakis, 2014) and ExaBayes v1.4.1 (Aberer et al., 2014), respectively. For both analyses we partitioned the dataset using a development version of PartitionFinder (PF), which implements a “kmeans” clustering algorithm to group similarly evolving sites together (Frandsen et al. 2015). This approach, developed specifically for non-coding loci like UCEs, identifies data subsets without user pre-partitioning. For the ML search we used the rapid bootstrapping plus best-tree search algorithm (“-f a” option, 100 bootstrap replicates) and we used GTR+Γ as the model of sequence evolution (for both best-tree and bootstrap searches). For the BI search we executed two independent runs, each with four coupled chains (one cold and three heated chains). We linked branch lengths across partitions and we ran each partitioned search for one million generations. We assessed burn-in, convergence among runs, and run performance by examining parameter files with the program Tracer v1.6.0 (Rambaut et al., 2014). We computed consensus trees using the *consense* utility, which comes as part of ExaBayes.

To assess the possible influence of gene-tree conflict we performed species-tree estimation using the summary method implemented in ASTRAL v4.8.0 (Mirarab et al. 2014). To reduce error from loci with low information content we employed weighted statistical binning, which bins loci together based on shared statistical properties and then weights bins by the number of included loci (Bayzid et al., 2015). To carry out the analysis, we first used RAxML to generate ML gene trees for all loci. We then input the gene trees into the statistical binning pipeline using a support threshold of 75 (recommend for data sets with < 1000 loci). This grouped genes into 168 bins, comprising 3 bins of 3 loci, 132 bins of 4 loci, and 33 bins of 5 loci. After binning we concatenated the genes into supergenes and used RAxML to infer supergene trees with bootstrap support (200 reps). We then input the resulting best trees, weighted by gene number, and the bootstrap trees, into ASTRAL and conducted a species tree analysis with 100 multi-locus bootstrap replicates (Seo, 2008).

### Divergence Dating

To generate a time tree for the evolution of stenammine ants we estimated divergence dates using the program BEAST v1.8 (Drummond et al., 2012). Due to computational challenges arising from having both a large number of taxa and a large amount of sequence data, we developed a two-part strategy to make the analyses feasible: (1) we input a starting tree (all nodes constrained) and turned off tree-search operators and (2) we input a subset of the sequence dataset rather than the entire concatenated matrix. For the input tree we used the best tree generated from the partitioned RAxML search and, for the data, we selected the 25 loci that had the highest gene tree bootstrap scores. The selected loci were concatenated together and analyzed without partitioning. We performed a total of four independent runs in BEAST, each for 100 million generations, sampling every one thousand generations. We also performed one run in which the data were removed so that the MCMC search sampled from the prior distribution only. For the clock model we selected uncorrelated lognormal, for the substitution model we used GTR+Γ, and for the tree prior model we used birth-death. For the ucld.mean prior we used an exponential distribution with mean 10.0.

To calibrate the analysis we used two secondary calibrations taken from Ward et al. (2015). These were used rather than using fossil calibrations because most of the fossils could not be placed within subgroups of Stenammini with high confidence. We applied one secondary calibration to the root node (crown-group Myrmicinae) and used a normal prior with mean 98.6 Ma and standard deviation 7 Ma. We applied the other calibration to crown-group Stenammini and used a normal prior with mean 52.9 Ma and a standard deviation of 4 Ma.

To evaluate the performance of BEAST runs we inspected run parameters using Tracer v1.6 (Rambaut et al. 2014). We looked for ESS values above 100 for all parameters and for convergence among runs. After deciding on an appropriate burn-in, we combined trees using *LogCombiner*, and generated a chronogram using *TreeAnnotator* (both programs included with the BEAST package).

### Biogeographic Reconstruction

We inferred the biogeographic history of Stenammini using the dispersal-extinction-cladogenesis model (DEC; Ree & Smith, 2008) as implemented in the program RASP (Yu et al., 2015). For input we used the consensus tree and 500 post-burnin trees from the BEAST divergence dating analysis. Using the R package APE (Paradis et al., 2004), we pruned all outgroup taxa from input trees, leaving only crown-group Stenammini. We coded terminal taxa using the six classic biogeographic realms (Olson et al., 2001; see Fig. 4c for coding): Nearctic (A), Neotropical (B), Palearctic (C), Afrotropical (D), Indomalayan (E), and Australasian (F). Based on the observation that most extant species occupy only one area, we set max areas to two regions. We excluded several implausible ancestral areas (AD, AE, AF, BC, BD, BE, BF, CF, CG, DE, DF) and reduced dispersal probabilities to 0 for several region pairs (AD, AE, AF, BC, BE) and to 0.5 for several others (BD, BF, DF). Given the limited amount of change in the position of continents during the timing of Stenammini evolution (< 55 Ma, see results below), we used a single time slice for the analysis. To take into account branch-length variation we performed a Bayes-DEC analysis in RASP using 100 post-burnin trees randomly selected from the pool of 500. RASP performs the DEC analysis on all 100 trees and plots the combined results on the consensus tree.

### Habitat Reconstruction

To gain further insight into niche evolution in Stenammini we inferred ancestral habitats using the DEC model in RASP (i.e., the same approach used for biogeographic reconstruction). All terminal taxa were coded as occupying one or a combination of the following habitats (see Fig. 4c for coding): temperate wet (A), temperate dry (B), tropical wet (C), or tropical dry (D). The implausible habitat combinations AD and BC were excluded as possible ancestral states and dispersal between these excluded habitat pairs was set to 0.5. For extratropical taxa we considered nesting biology when coding species as being either wet or dry adapted. Some *Veromessor* species, for example, occur in areas in the western USA that receive periodic rainfall, but they nest in dry, open habitats. We coded these taxa as B. Other species occur in the same general region as *Veromessor*, but are only active in moist leaf litter (e.g., *Stenamma*). We coded these taxa as A. Several extratropical taxa with uncertain or intermediate nesting habitats were coded as AB.

### Diversification Rates

We investigated diversification dynamics in Stenammini using two main approaches. First, we tested for significant shifts in diversification rate across the entire tree using the R package TreePar (Stadler, 2011). TreePar is a ML-based program that allows for non-constant diversification rates and incomplete taxon sampling. It fits a birth-death model to dated trees and allows rates to change as a function of time. We ran the program using the “bd.shifts.optim” function and used the dated consensus tree from BEAST as the input tree (outgroups pruned). We set extant diversity at 22% to account for incomplete species sampling (92/416 species sampled, including three undescribed species) and we set the program to test for a maximum of five rate shifts at 0.1 My intervals ranging from zero to 53 Ma. We selected the best-fitting model by performing likelihood-ratio tests with significance assessed by P-values below 0.05.

After assessing tree-wide rate shifts, we tested for rate shifts among lineages using the Bayesian program BAMM v2.5 (Bayesian Analysis of Macroevolutionary Mixtures; Rabosky et al., 2013, 2014a; Rabosky, 2014; Shi & Rabosky, 2015) and the accompanying R package BAMMtools (Rabosky et al., 2014b). BAMM is capable of modeling complex diversification dynamics by exploring many models simultaneously using reversible jump Markov Chain Monte Carlo (rjMCMC). A major advantage over likelihood-based approaches is that this method allows one to assess support for different rate-shift configurations using posterior probabilities and Bayes Factors. It also allows for incomplete taxon sampling in a way that does not require collapsing or pruning branches in the input tree.

For input into BAMM we used the BEAST consensus chronogram with outgroups pruned. To account for non-random, incomplete taxon sampling we input a sampling probability file, in which we assigned all terminals to clades roughly corresponding to genera (*Aphaenogaster*, “Deromyrma”, “Goniomma”, *Messor*, *Novomessor*, *Stenamma, Veromessor*; see Fig. 3) and we gave each clade a sampling probability based on the number of species sampled and the number of extant species in that clade (0.10, 0.46, 0.40, 0.05, 1.00, 0.45, and 0.89, respectively). To select appropriate priors for the BAMM analysis, we used the “setBAMMpriors” function in BAMMtools on the input time tree. Using the prior settings, we then ran the BAMM analysis with four chains for 2,000,000 generations, sampling event data every 1,000 generations. After the analysis, we assessed burnin, run convergence, and ESS values for all parameters using the R package CODA (Plummer et al., 2006). We then explored the post-burnin output using BAMMtools and selected the best rate-shift configuration by assessing posterior probabilities and Bayes Factors.

To visualize lineage accumulation through time, we used the APE package in R to generate a lineage-through-time plot from the BEAST consensus tree and 500 post-burnin trees

## RESULTS

### Diversity Gradient

We calculated latitudinal midpoints for all 413 extant species currently placed in the tribe Stenammini (Fig. 1). Using a cutoff for the tropics at 23.5°N/S latitude, we found that 340 species are extratropical and only 73 species are tropical. The tribe ranges from roughly 30°S to 50°N and has an obvious diversity peak in the northern hemisphere between 35°N to 40°N latitude. The diversity distribution is strongly left-skewed, with richness dropping below ten species from 10°N to 35°S (binned at 5° intervals). This distribution is concordant with Stenammini having an inverse LDG, and thus being a contrarian clade.

**Figure 1.**
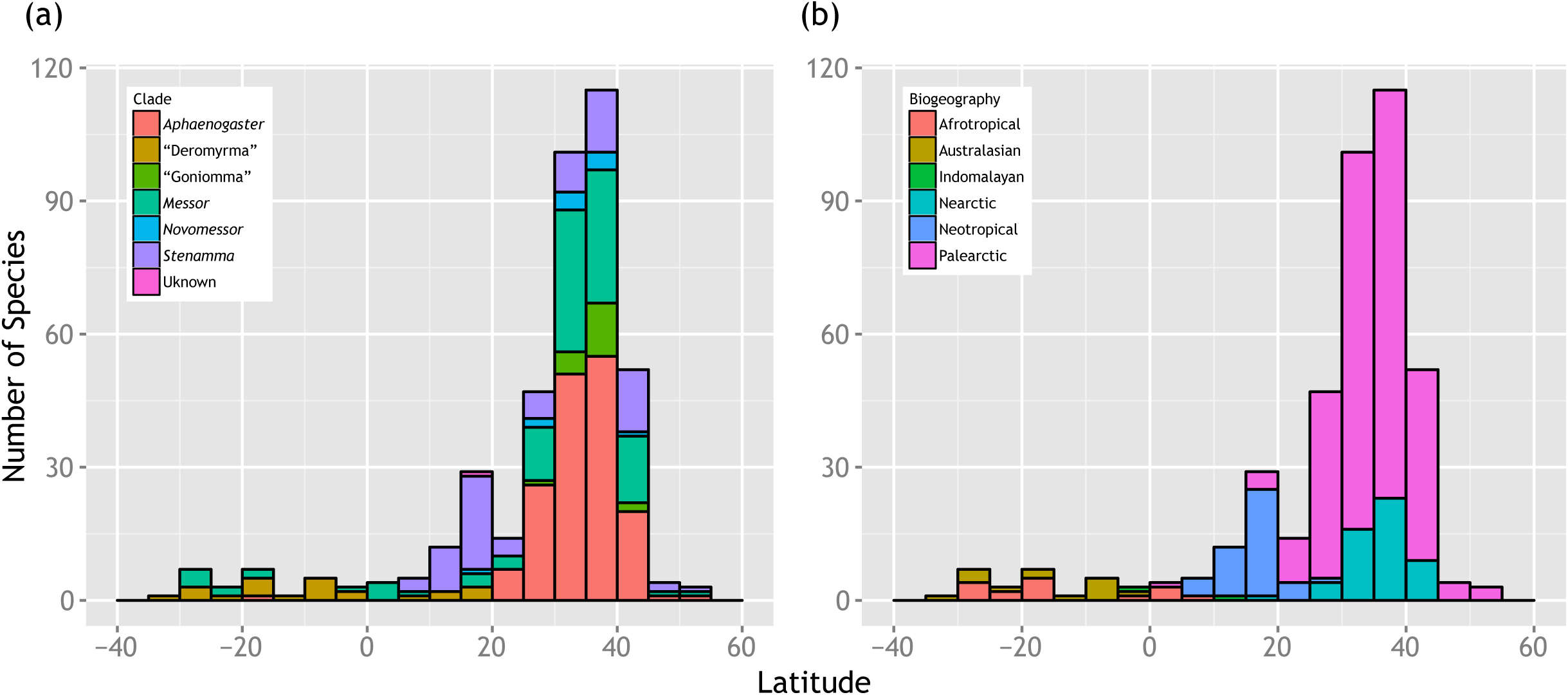
Midpoint latitude of all extant species of Stenammini binned into 5° intervals. The plot shows a strong richness peak in the northern temperate zone. (a) Bars colored by major clade. The “Unknown” category corresponds to the enigmatic species *Aphaenogaster relicta*. (b) Bars colored by biogeography.

### UCE Phylogeny

Following sequencing, assembly, and the matching of probes to contigs, we recovered an average of 987 UCE contigs per sample, with a mean contig length of 909 bp and an average coverage score of 85X (see Appendix S3). We evaluated the results of filtering loci for taxon completeness and selected the 90% filtered matrix (*Filter90*) as the primary locus set for analysis (see Appendix S4 for filtering results). Following sequence alignment, the *Filter90* locus set included 702 loci and had an average locus length of 849 bp, resulting in a concatenated data set of 595,931 bp, of which 198,260 sites were informative. All sequences have been submitted to the NCBI Sequence Read Archive (upon acceptance) and the *Filter90* concatenated data set has been submitted to TreeBASE (upon acceptance)

The partitioned ML and BI analyses of the concatenated data set resulted in identical topologies, with most nodes receiving maximum support (Fig. 2 and Appendix S5). The few nodes with less than complete support were restricted to the shallower splits in the tree. The ST analysis produced a mostly congruent tree (Appendix S5), with one major difference involving the position of the genus *Messor* (discussed below).

**Figure 2.**
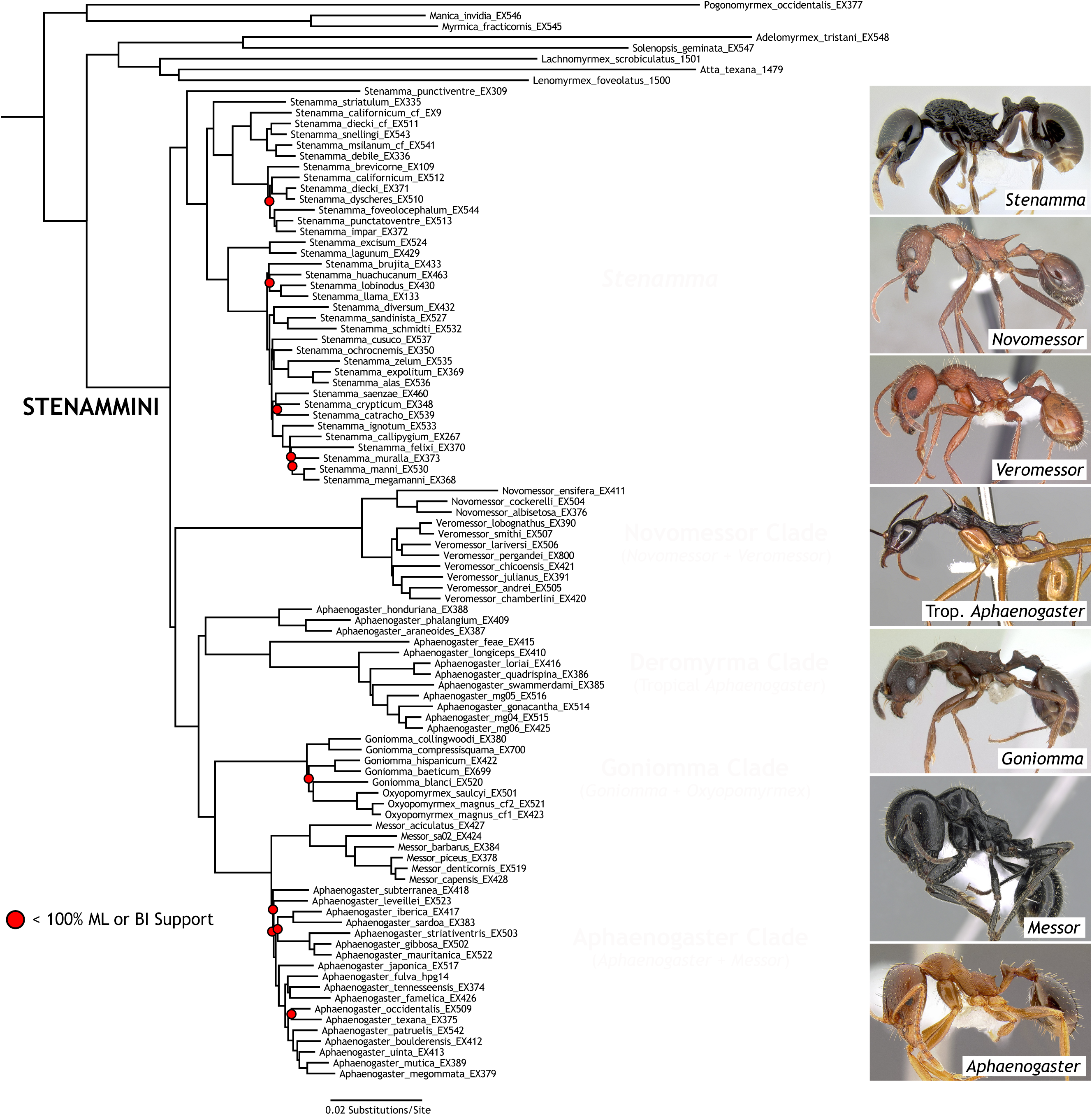
Best maximum likelihood tree depicting relationships among species of Stenammini. Red dots indicate nodes that received less than 100% bootstrap support (RAxML) or posterior probability (ExaBayes). All other nodes received maximum support. Major lineages are highlighted and named as follows: *Stenamma*, “Novomessor” clade, “Deromyrma” clade, “Goniomma” clade, and “Aphaenogaster” clade. Ant images from www.antweb.org.

We recovered the tribe Stenammini as monophyletic and comprising five major clades: (1) the genus *Stenamma*, (2) the genera *Novomessor* + *Veromessor* (the “Novomessor” clade), (3) a group of tropical “Aphaenogaster” (the “Deromyrma” clade), (4) the genera *Goniomma* plus *Oxyopomyrmex* (the “Goniomma” clade), and (5) the genus *Messor* plus all Holarctic members of *Aphaenogaster* (the “Aphaenogaster” clade). The distributions of major groups are provided in Fig. 3.

**Figure 3.**
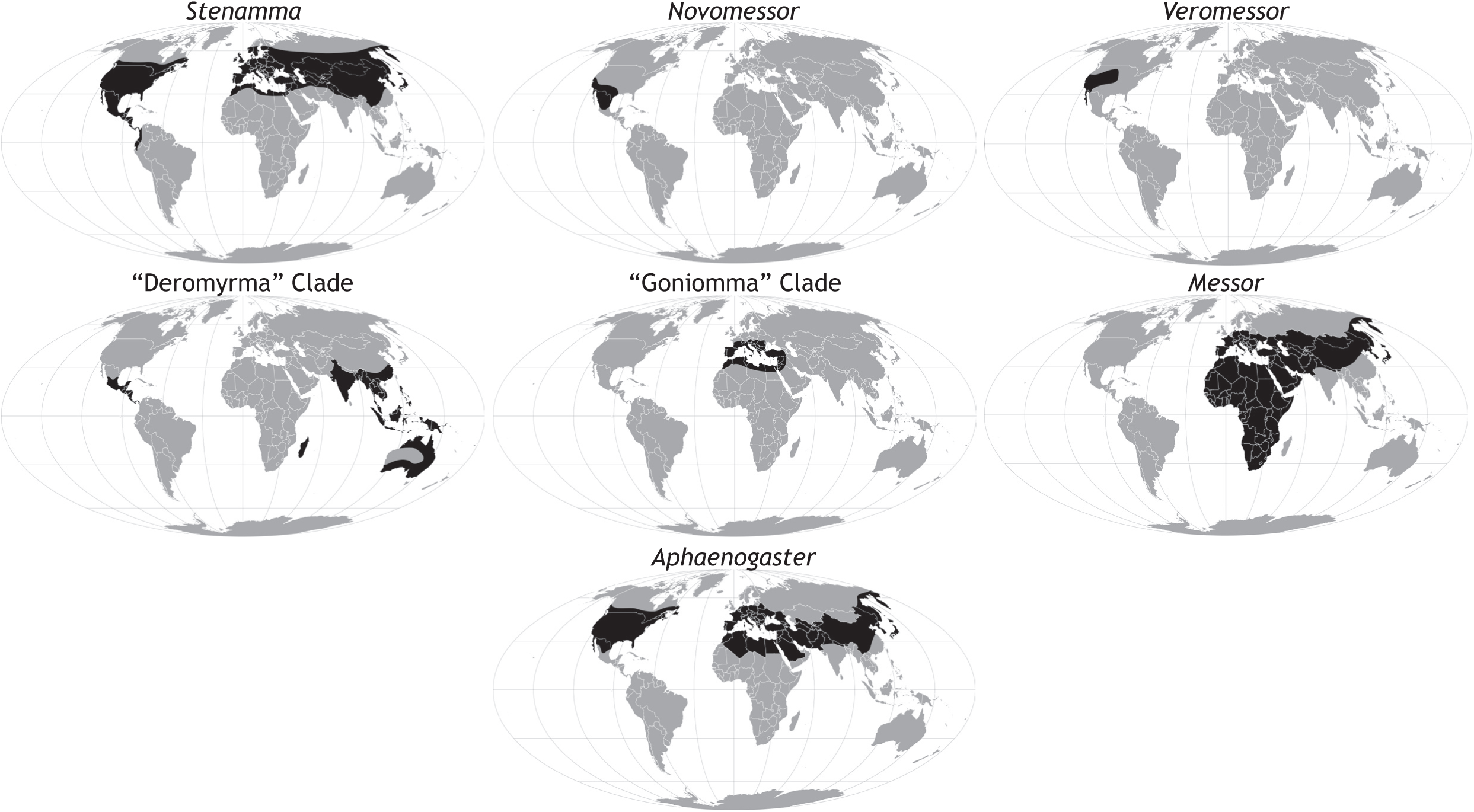
Distribution maps for major lineages of Stenammini. Maps are Mollweide equal-area projections.

The genus *Stenamma* was recovered as sister to the rest of the stenammine clades and within *Stenamma*, the species *S. punctiventre*, a relict species form Morocco, was found to be sister to all remaining species, which themselves separate into two major clades that have previously been referred to as the “Holarctic” and “Middle American” clades (Branstetter, 2012). These two clades are nearly geographically isolated, overlapping only slightly in the southwestern USA.

Outside of *Stenamma*, the “Novomessor” clade was recovered as sister to all remaining clades, and the genera *Novomessor* and *Veromessor* were each found to be monophyletic. The “Deromyrma” clade was recovered as sister to the “Goniomma” clade plus the “Aphaenogaster” clade. Unexpectedly, the “Deromyrma” clade was found to include all tropical species of the genus *Aphaenogaster*, and these clustered by region into New World and Old World clades. Within the “Goniomma” clade the genus *Goniomma* was found to be paraphyletic with respect to *Oxyopomyrmex*.

In the “Aphaenogaster” clade we obtained conflicting results with respect to the position of the genus *Messor*. In both the partitioned ML and BI analyses *Messor* is monophyletic and sister to a monophyletic *Aphaenogaster*. However, in the ST analysis the *Aphaenogaster* species *A. subterranean* and *A. leveillei* were found to be sister to *Messor*, making *Aphaenogaster* paraphyletic. In the preliminary ML analyses in which we assessed different levels of missing data, we also recovered *Aphaenogaster* paraphyly (see Appendix S4). Uncertainty in *Aphaenogaster* monophyly is also indicated in the partitioned ML and BI trees by low support and a very short branch subtending the clade (Fig. 2).

### Historical Biogeography

The following focuses on key nodes and combines the results of the BEAST divergence dating analysis and the S-DEC biogeographic reconstruction. Dates are presented as median ages and ancestral areas are presented as the most likely state, except in cases in which different inferred areas have nearly the same probability. Results are summarized in Fig. 4c, Table 1, and Appendix S6.

**Table 1.**
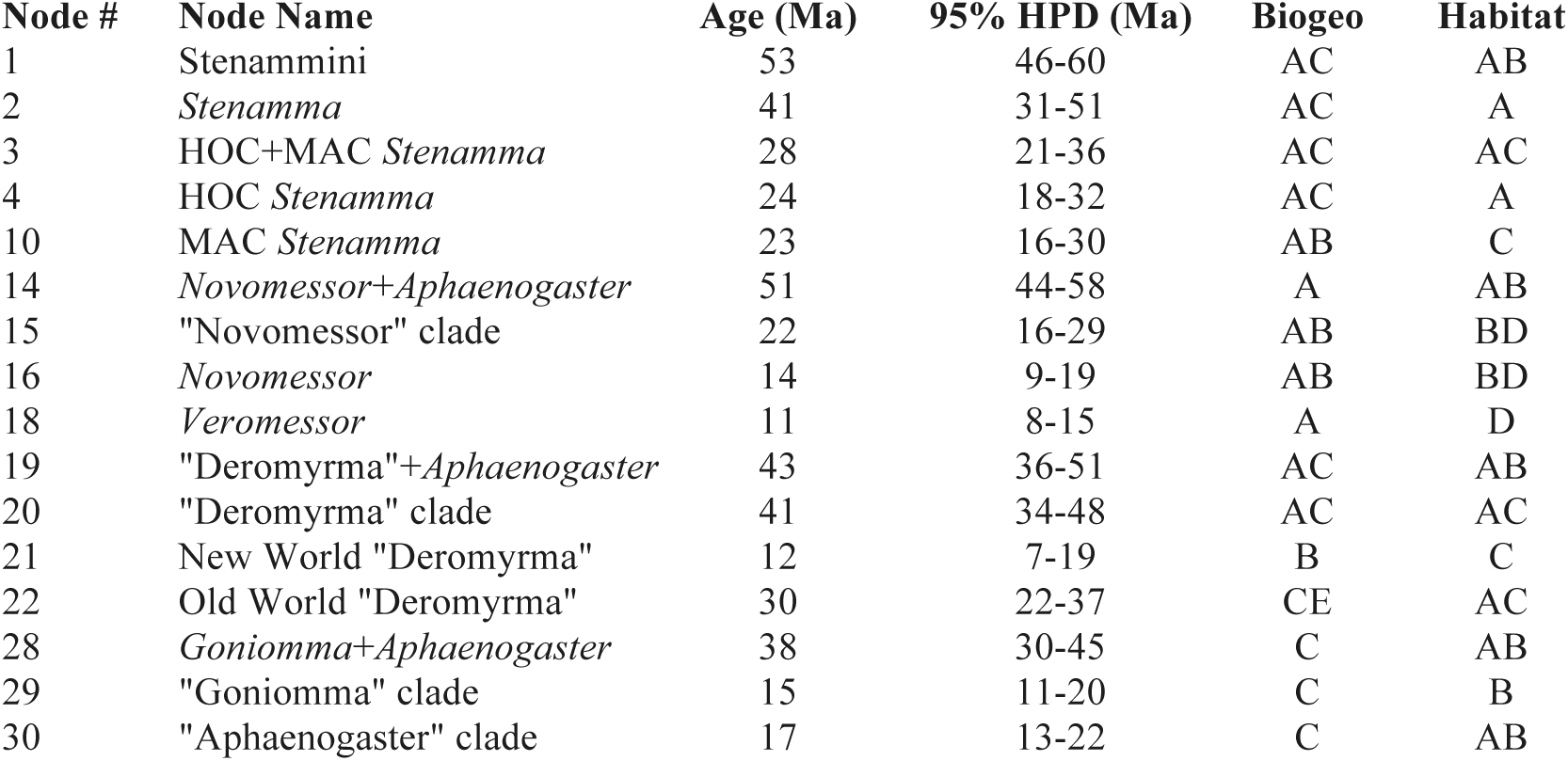
Results from divergence dating, biogeography, and habitat reconstruction. Node numbers correspond to the numbers in Fig. 4c. Biogeographic areas: Nearctic (A), Neotropics (B), Palearctic (C), Afrotropics (D), Indomalaya (E), Australasia (F). Habitats: temperate wet (A), temperate dry (B), tropical wet (A), tropical dry (B).

| Node # | Node Name | Age (Ma) | 95% HPD (Ma) | Biogeo | Habitat |
| --- | --- | --- | --- | --- | --- |
| 1 | Stenammini | 53 | 46-60 | AC | AB |
| 2 | *Stenamma* | 41 | 31-51 | AC | A |
| 3 | HOC+MAC *Stenamma* | 28 | 21-36 | AC | AC |
| 4 | HOC *Stenamma* | 24 | 18-32 | AC | A |
| 10 | MAC *Stenamma* | 23 | 16-30 | AB | C |
| 14 | *Novomessor*+*Aphaenogaster* | 51 | 44-58 | A | AB |
| 15 | "Novomessor" clade | 22 | 16-29 | AB | BD |
| 16 | *Novomessor* | 14 | 9-19 | AB | BD |
| 18 | *Veromessor* | 11 | 8-15 | A | D |
| 19 | "Deromyrma"+*Aphaenogaster* | 43 | 36-51 | AC | AB |
| 20 | "Deromyrma" clade | 41 | 34-48 | AC | AC |
| 21 | New World "Deromyrma" | 12 | 7-19 | B | C |
| 22 | Old World "Deromyrma" | 30 | 22-37 | CE | AC |
| 28 | *Goniomma*+*Aphaenogaster* | 38 | 30-45 | C | AB |
| 29 | "Goniomma" clade | 15 | 11-20 | C | B |
| 30 | "Aphaenogaster" clade | 17 | 13-22 | C | AB |

**Figure 4.**
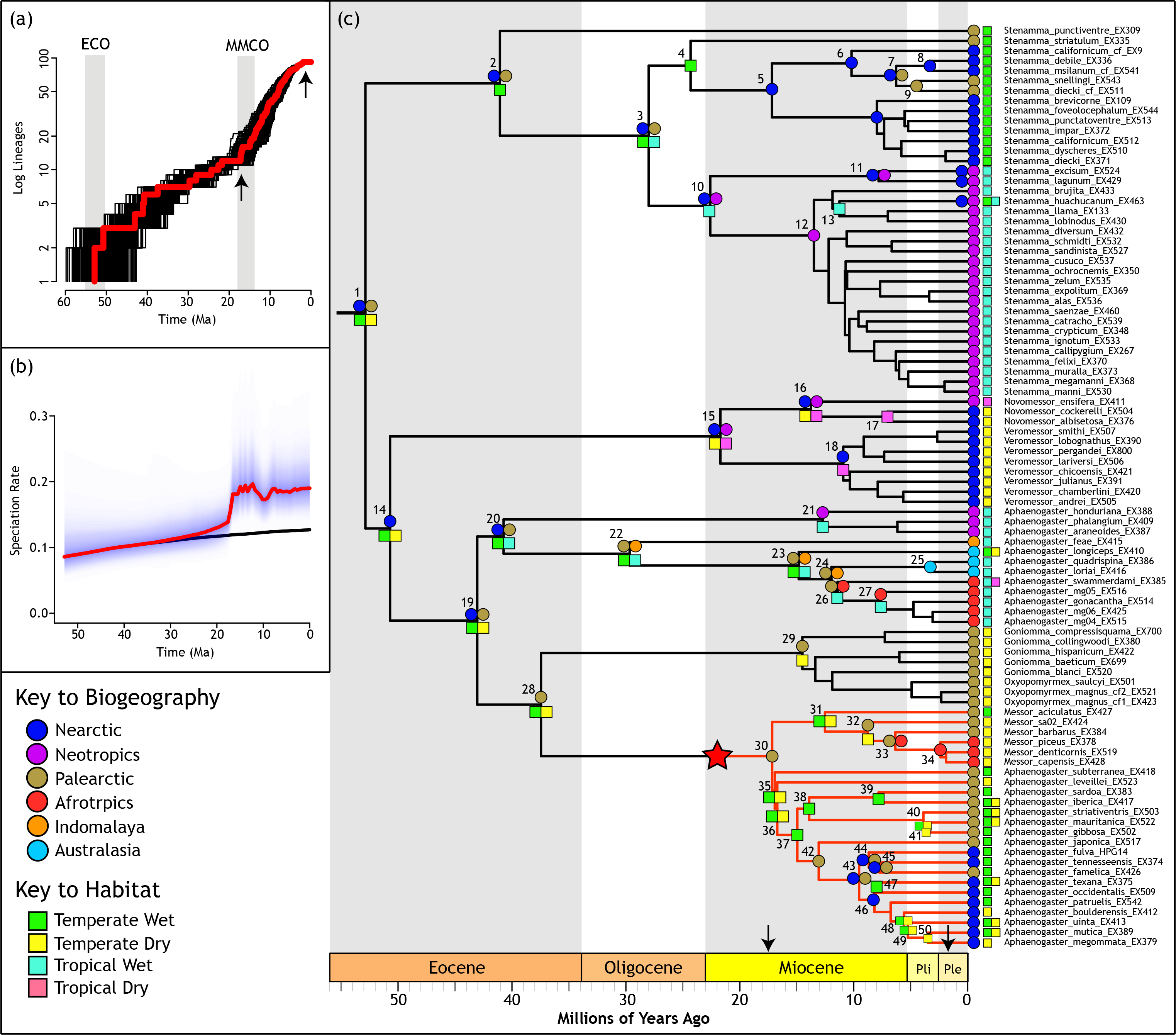
Results from divergence dating, ancestral area inference, habitat reconstruction, and diversification rate tests. (a) Lineage-through-time plot, with TreePar tree-wide rate shifts marked by arrows and the Eocene Climatic Optimum (ECO) and Mid-Miocene Climatic Optimum (MMCO) marked with grey bars. (b) BAMM speciation-rate-through-time plot showing the overall rate (red line) and the rate with the “Aphaenogaster” clade excluded. (c) Maximum clade credibility tree from divergence-dating analysis with BEAST. The red star and red branches indicate the rate shift inferred by BAMM. The black arrows along the geological timeline mark the tree-wide rate shifts inferred by TreePar. Circles and squares on nodes represent inferred ancestral areas and ancestral habitats, respectively. Only key nodes are marked.

The tribe Stenammini evolved between 89-52 Ma (age between stem and crown groups), with the crown group originating in the Holarctic region. Among the major clades, most have an origin in the northern temperate zone. The genus *Stenamma* originated in the Holarctic 41 Ma. The “Goniomma” and “Aphaenogaster” clades both originated in the Palearctic region 15 Ma and 17 Ma, respectively. Unexpectedly, the “Deromyrma” clade, which includes mostly tropical taxa (a few species occur in temperate Australia), also was found to have a temperate origin, with the ancestor inhabiting the Holarctic 41 Ma. The only major clade that may not have had a temperate-only origin was the “Novomessor” clade. This group originated 22 Ma and was inferred to have either a Nearctic plus Neotropical origin (slightly higher likelihood) or a Nearctic-only origin. Within the “Novomessor” clade, the DEC model inferred a Nearctic plus Neotropical origin for *Novomessor* and a Nearctic-only origin for *Veromessor*, 14 Ma and 11 Ma, respectively.

Across the tribe there were a total of six separate dispersal events out of the extratropics and into the tropics. The ancestor of the Middle American clade of *Stenamma* dispersed into the tropics 28-23 Ma. Depending on the reconstruction, the ancestor of the “Novomessor” clade dispersed into the tropics between 50-22 Ma or the ancestor of *Novomessor* dispersed into the tropics between 22-14 Ma. The genus *Messor* dispersed into the Afrotropics more recently, 9-6 Ma. In the “Deromyrma” clade we inferred three separate dispersals into the tropics, once into the Neotropics 41-13 Ma, once into Indomalaya 41-30 Ma, and once into the Afrotropics 12-11 Ma.

Dispersals out of the tropics and between the Nearctic and Palearctic regions also occurred. We inferred three separate out-of-the-tropics dispersal events. The ancestor of *S. huachucanum* back-dispersed from the Neotropics to the Nearctic between 11 Ma and the present. This species has a Neotropical plus Nearctic distribution, reaching as far north as the sky islands of Arizona. In the genus *Messor*, dispersal into the Afrotropics included dispersal into the arid subtropics of South Africa between 7-2 Ma. Similarly, the Australian species *A. longiceps*, part of the “Deromyrma” clade, occurs in tropical and extratropical Australia, likely having dispersed out of the tropics between 15 Ma and the present. Dispersal from the Nearctic to the Palearctic occurred once in the Holarctic clade of *Stenamma*, most likely between 10-6 Ma. Finally, dispersal from the Palearctic into the Nearctic occurred in Holarctic *Aphaenogaster* between 13-10 Ma.

### Niche Reconstruction

We inferred ancestral habitats using the Bayes-DEC model in RASP (Fig. 4c, Table 1, and Appendix S6). Concordant with the biogeographic results, for most major clades we recovered temperate habitats for ancestral nodes. For crown-group Stenammini we inferred a temperate wet plus temperate dry ancestral habitat. For the genus *Stenamma* we inferred a temperate wet ancestor, with a shift into temperate wet plus tropical wet habitats in the ancestor of the Holarctic plus Middle American clades. The “Goniomma” clade had a temperate dry ancestor and the “Aphaenogaster” clade had a temperate wet plus temperate dry ancestor. Both the “Novomessor” and “Deromyrma” clades had temperate plus tropical ancestral habitats. In the “Novomessor” clade we inferred the habitat to be temperate dry plus tropical dry. In the “Deromyrma” clade we inferred the ancestor to have a temperate wet plus tropical wet ancestral habitat, with most species shifting into a tropical wet habitat only.

### Diversification Dynamics

Analysis with the program TreePar, which tests for diversification rate shifts across a tree, found two rate shifts during stenammine evolution: one increase in net diversification rate at 17.2 Ma during the Miocene (shift from 0.046 to 0.227 species Myr^−1^), and one decrease in rate at 1.9 Ma in the Pleistocene (shift from 0.227 to 001 species Myr^−1^; Fig. 4ac & Appendix S7).

Complementing the TreePar results, diversification analysis with BAMM, which tests for rate shifts among lineages, found high support for an increase in diversification rate in the branch leading to the “Aphaenogaster” clade (Fig. 4c). The BAMM run produced posterior probabilities for seven rate-shift models. Of these, the single rate-shift model received the highest posterior probability (74%) and was favored by Bayes Factor comparison as compared to the zero rate shift model (BF score of 2,666) and all other shift models (see Appendix S7). The “Aphaenogaster” clade is dated to 17.2 Ma with the rate shift inferred to have occurred along the stem lineage several million years prior to the origin of the crown group. Examining speciation rates through time, the results show very similar, constant rates for all lineages except for the “Aphaenogaster” clade (Fig. 4b, Table 2). The average speciation rate for the entire tree was 0.174 species Myr^−1^ and the background rate (“Aphaenogaster” clade excluded) was 0.123 species Myr^−1^. The rate for just the “Aphaenogaster” clade was 0.363 species Myr^−1^.

**Table 2.** Average speciation and extinction rates estimated in BAMM for key groups of taxa. Node numbers correspond to the numbers in Fig. 4c. Note that rates are similar except for taxon groups that include the “Aphaenogaster” clade, which has a significantly increased rate. HOC: Holarctic Clade. MAC: Middle American Clade.

| Node # | Lineage | $\lambda$ Mean | $\mu$ Mean |
| --- | --- | --- | --- |
| 1 | all Stenammini | 0.1744 | 0.0430 |
| 1 | no "Aphaenogaster" clade | 0.1226 | 0.0219 |
| 30 | "Aphaenogaster" clade | 0.3627 | 0.1199 |
| 41 | *Stenamma* | 0.1241 | 0.0219 |
| 4 | HOC *Stenamma* | 0.1255 | 0.0217 |
| 10 | MAC *Stenamma* | 0.1268 | 0.0217 |
| 16 | "Novomessor" clade | 0.1214 | 0.0219 |
| 20 | "Deromyrma" clade | 0.1213 | 0.0219 |
| 29 | "Goniomma" clade | 0.1235 | 0.0218 |

## DISCUSSION

The current latitudinal diversity gradient, whereby species richness increases from pole to equator, is the predominant diversity pattern found in most major organismal lineages. An increasing number of groups, however, are known to exhibit non-standard gradients. In this study we have added the ant tribe Stenammini, a large taxon with over 400 species, to this growing list of contrarian clades. Our results show that the tribe exhibits a strong inverse diversity gradient, with the majority of taxa occurring in the northern temperate zone. Following this observation we used phylogenomic techniques to establish a robust phylogeny for the tribe and we used the phylogeny to evaluate alternative hypotheses that might explain the group’s non-standard diversity pattern.

### The latitudinal diversity gradient

Using midpoint latitudes for all extant stenammine species, we demonstrated that the tribe has a strong inverse diversity gradient, with the majority of species inhabiting the northern temperate zone between 30-45°N latitude (Fig. 1). Separating results by clade (Fig. 1a) and biogeographic region (Fig. 1b), the diversity peak is mainly attributable to *Aphaenogaster* and *Messor* species in the Palearctic. Could this pattern be driven by a taxonomic or sampling bias? We strongly believe that the answer is no. In recent years there has been extensive ant sampling by the authors and colleagues in Central America, Africa, Madagascar, and Southeast Asia, and few additional tropical morphospecies have been identified. Further, the genus *Stenamma*, which has the highest tropical diversity among genera in the clade, was recently revised (Branstetter, 2013) and we accounted for all new species in our analysis. In contrast to tropical diversity, there have been several recent taxonomic revisions focusing on Palearctic taxa and these studies have all added species to overall diversity (Terayama, 2009; Boer, 2013; Borowiec & Salata, 2013, 2014; Kiran et al., 2013; Salata & Borowiec, 2015). Furthermore, it is likely that many new Palearctic species await discovery in poorly sampled regions, such as the southern Mediterranean and Middle East.

A growing number of lineages from both the marine and terrestrial realms have been identified as contrarian clades with extratropical peaks in species richness (Owen & Owen, 1974; Dixon et al., 1987; Chown et al., 1998; Stephens & Wiens, 2003; Smith et al., 2005; Krug et al., 2007; Pyron & Burbrink, 2009; Rivadeneira et al., 2015). When considering these studies, it is important to distinguish between clade-based and assemblage-based approaches. The clade-based studies have taken into account species richness patterns for an entire monophyletic group. The assemblage-based studies, however, investigated communities of species and are often spatially restricted. Although the standard LDG might be considered an assemblage-based phenomenon, we argue that identifying and studying contrarian clades helps to explain diversity gradients broadly, especially when the clades of interest are diverse and widespread. In our study we have taken a clade-based approach and have shown that the ant tribe Stenammini is a diverse lineage that defies the standard LDG. Of the available clade-based studies, in which a phylogenetic approach has been used to examine an inverse LDG, ours represents the largest to date in terms of number of species considered.

An interesting characteristic of the inverse gradient observed in Stenammini is its skewed distribution. Even though the group extends into the southern temperate zone, the diversity peak is only found in the northern hemisphere. This pattern of a northern temperate species peak has been observed in lampropeltinine snakes (Pyron & Burbrink, 2009), emydid turtles (Stephens & Wiens, 2003), and several clades of hylid frogs (Smith et al., 2005). Salamanders also show this pattern at the family level, but not at the species level (Wiens, 2007). In contrast, marine bivalves (Krug et al., 2007) and aphids (Dixon et al., 1987) have diversity peaks in both the northern and southern hemispheres. Currently there are no clade-based studies that have shown a skewed distribution in which diversity peaks in the southern temperate zone. A number of studies have shown an inverse diversity gradient along the coast of South America, but these studies have focused on species assemblages rather than clades (Rivadeneira et al., 2011). Other groups with temperate richness peaks but uncertain symmetry include pelagic seabirds (Chown et al., 1998) and ichneumonid wasps (Owen & Owen, 1974). As discussed below this skewed distribution of Stenammini is likely due its origin in the northern hemisphere.

### Evolution of Stenammini

Crown-group Stenammini originated in the Holarctic during the early Eocene approximately 53 Ma. This time period corresponds to the Eocene Climatic Optimum (ECO) when temperatures were on average 5-6°C warmer than today (Zachos et al., 2001) and broad-leaved evergreen forest and paratropical rainforest were greatly expanded (Graham, 1993, 2011). Fossil records indicate that broad-leaved, evergreen forest extended above 70°N latitude (Eberle & Greenwood, 2012; Harrington et al., 2012) and that animals with present-day tropical affinities, such as the Arctic tapir, inhabited these forests (Eberle & Greenwood, 2012). It is also known that the vegetation in North America and Eurasia was more homogeneous during this time, likely due to the lack of polar glaciation and the existence of two major land bridges, the Bering Land Bridge (75°N) and the North Atlantic Land Bridge (45-65°N) (Wing, 1987). Stenammini, therefore, originated in the Holarctic at a time when wet forest habitat dominated the northern hemisphere and significant connections between northern continents existed, facilitating dispersal. Our inferred ancestral habitat for the tribe of temperate wet and temperate dry is in line with the idea that the ancestral stenammine species could have existed throughout the Holarctic in both the dominant forest habitat and the more limited deciduous and dry habitats.

After the ECO there was a gradual decrease in global temperature culminating in a sharp decrease 35 Ma at the Eocene-Oligocene boundary. During the cooling, the evergreen forests retracted from their northern maxima, and more seasonal habitats, including deciduous forest, expanded (Wing, 1987; Graham, 2011). This period of time is important because many modern plant and animal lineages originated (Wing, 1987; Pellmyr & Leebens-Mack, 1999; Bininda-emonds et al., 2007; Jaramillo et al., 2010). For plants the homogeneous nature of the vegetation between North American and Eurasia began to disappear, suggesting that dispersal via the northern land bridges became increasingly less likely, especially for species adapted to warm, wet environments. This observation that wet-adapted species were once widespread across Eurasia, but then had to retract into tropical refugia has become known as the Boreotropics hypothesis and is commonly proposed in phytogeography to explain taxa with disjunct distributions between the Old World and New World tropics (Wolfe, 1975; Tiffney, Bruce, 1985; Lavin & Luckow, 1993; Donoghue, 2008). We believe this scenario is important in considering the evolution of Stenammini, especially for the “Deromyrma” clade.

The “Deromyrma” clade can be aptly described as a tropical group even though a few species extend into the Australian temperate zone. Our phylogenetic results show that the clade originated ~41 Ma and bifurcated into New and Old World lineages. Contrary to expectations, we found that the clade originated in the Holarctic and dispersed into the tropics three times, once into Mesoamerica, once into Indomalaya, and once into the Afrotropics. Our biogeographic reconstruction is completely concordant with the Boreotropics hypothesis. The ancestor of the “Deromyrma” clade was widespread and adapted to wet evergreen forest habitat. When the wet forest contracted and no longer extended across the northern land bridges, “Deromyrma” separated vicariantly into the New and Old World lineages. Both of these lineages then followed the retraction of wet forest into the tropics, where they have persisted to the present. This scenario is concordant with the fossil record, which includes a species from Dominican Amber (*Aphaenogaster amphiocenica*) that has been suggested to be closely related to tropical *Aphaenogaster* (De Andrade, 1995)

In the Old World tropics the “Deromyrma” clade has penetrated into Indomalaya, Australasia, and Madagascar (Afrotropics). How the clade arrived in Madagascar is interesting to consider, since it is not known from mainland Africa. Our biogeographic reconstruction suggests that the clade dispersed into the Afrotropics from the Palearctic rather than from Indomalaya or Australasia. This scenario suggests dispersal through Africa to reach Madagascar with later extinction from the mainland. A “through-Africa” hypothesis is supported by the fact that most extant Malagasy taxa have African origins (Yoder & Nowak, 2006) and rainforest was once more extensive in Africa during the Eocene (Burgess et al., 1998; Couvreur et al., 2008).

Global cooling explains another intriguing diversification event within the Stenammini. The ant genus *Stenamma* includes three clades, the relict Palearctic taxon *S. punctiventre*, a Holarctic clade, and a Middle American clade. The species *S. punctiventre* is included in a phylogeny for the first time here. Unexpectedly, the species was placed as sister to all remaining *Stenamma*, pushing the age of the crown group back to 41 Ma. Previous work inferred the origin of *Stenamma* to be Nearctic (Branstetter, 2012), but with *S. punctiventre* included, we inferred the origin of the genus to be Holarctic. The two major clades in the genus each contain about 40 species, with the Holarctic clade including a mix of Nearctic and Palearctic lineages and the Middle American clade including predominately Central American species, with only a couple of species reaching northern Mexico/southern U.S.A. and a few reaching northern South America. The timing of the split for these two clades is ~28 Ma. This date overlaps closely with the sharp cooling event at the Eocene-Oligocene boundary, which caused the mesic forests that once stretched between North and Central America to become fragmented due to aridification in the southwestern USA and northern Mexico (Wing, 1987). Given that *Stenamma* is generally associated with wet forest leaf litter, this aridification event is likely what caused the nearly complete split between northern and southern clades. More recently, the Middle American clade back-dispersed to the Nearctic via montane sky islands (e.g., *Stenamma huachucanum*).

The strong association of Stenammini with the northern temperate zone is further exemplified by the “Goniomma” and “Aphaenogaster” clades having ancestral areas in the Palearctic region. The “Goniomma” clade originated ~15 Ma and is completely restricted to the Mediterranean region. The “Aphaenogaster” clade is about the same age (17 Ma), but has spread more widely across the globe. It is distributed across the Holarctic and with the genus *Messor* has penetrated into arid regions in Africa, extending all the way down into South Africa.

The only major clade that does not have a completely north-temperate origin is the “Novomessor” clade. This group is arid-adapted and likely evolved in the Nearctic plus Neotropics or just the Nearctic with later dispersal into the Neotropics. The age of the clade is 22 Ma and it is split into the two monophyletic genera *Novomessor* and *Veromessor*. The former is distributed more southerly into Mexico, whereas the latter is more diverse in the western USA. It is possible that these two genera separated into northern and southern lineages during periods of increased aridification, similar to the pattern observed in *Stenamma*.

The results of the diversification analysis provide a compelling scenario for how Stenammini reached its current species-richness levels. Considering the across-tree analysis, we inferred an increase in diversification rate at 17.2 Ma and a slowdown at 1.9 Ma (Fig. 4a-c). This time period includes the origin of all of the more diverse clades in the tribe and it corresponds to the Mid-Miocene Climatic Optimum (MMCO) and a major decrease in global temperature that has continued to the present. Thus, while most higher-level diversification in the tribe is linked to global cooling after the ECO, most species-level diversity is correlated with cooling following the MMCO. It is very likely that the fragmentation of habitats caused by increased aridification during this time contributed to allopatric diversification in the tribe. The other important result is that among lineages, there was a significant increase in diversification rate leading to the “Aphaenogaster” clade. The importance of this rate shift to the overall diversification dynamics is obvious when the “Aphaenogaster” clade is excluded and only the background rate is examined (Fig. 4b, black line). This result suggests that most stenammine lineages have been evolving at a constant rate, but that the “Aphaenogaster” clade experienced a significant rate increase starting just prior to 17.2 Ma.

Why did the “Aphaenogaster” clade diversify significantly faster than the other lineages? We believe the answer can be linked to the group’s ecological diversity and its ties to the Mediterranean region. The clade includes two major genera, *Messor* and *Aphaenogaster*. The former is a specialist seed-harvester that is well adapted to desert and grassland habitats. These adaptations include size polymorphism, with the soldiers having large, muscular heads for eating seeds, and long beard-like hairs underneath the head, which are used for transporting excavated sand. *Messor* is currently widespread, occurring throughout the Palearctic and into arid Africa. The genus *Aphaenogaster* is more varied in its habits, but tends to be a medium-sized epigeic ant that is omnivorous. It does well in temperate forests and seasonally dry areas around the Mediterranean. Given the ability of both genera to do well in seasonally dry areas, increasing aridification over the last 20 Ma may have promoted diversification by fragmenting forested habitats and expanding arid environments.

The overall pattern of evolution in Stenammini suggests that it has been most successful in temperate habitats. All the major clades evolved in one or both of the Holarctic regions and the majority of the species diversity is found there. Even though the clade has dispersed to the tropics at least six times, the majority of those tropical lineages have failed to diversify greatly. The only exception to this is the ant genus *Stenamma*, which includes a lineage that has radiated into over 40 species in Central America. This lineage, however, does not exhibit a significantly higher diversification rate as compared to its Holarctic sister clade and it shows a clear connection to its temperate past: it is a montane specialist with greatest diversity and abundance in wet forest above 1,000 m. This pattern contrasts with ants generally, which show greater diversity and abundance in the tropical lowlands.

### Evaluating alternative LDG hypotheses

Using our results we can differentiate among several competing hypotheses that might explain the inverse latitudinal gradient exhibited by Stenammini. These hypotheses are the time-for-speciation/center-of-origin (TFS/COO) hypothesis (Schluter, 1993; Wiens & Donoghue, 2004; Stevens, 2006), the niche conservatism hypothesis (Wiens & Donoghue, 2004; Wiens & Graham, 2005), and the diversification rate hypothesis (Rohde, 1992; Cardillo et al., 2005; Weir & Schluter, 2007). Given that Stenammini is most diverse in the northern temperate zone, the TFS/COO would predict that the clade originated in this region and thus has had more time to diversify there. This prediction is supported. The tribe originated in the Holarctic, as did most of the major clades. It is also supported by the fact that the Middle American and Holarctic clades of *Stenamma* have nearly the same number of species. These clades are the same age and thus should have about the same number of species under the TFS/COO.

The niche conservatism hypothesis predicts that niche evolution is rare and that the niche with the most species will be the ancestral niche. For Stenammini, this hypothesis is partly supported. We inferred the ancestral habitat to be temperate wet and temperate dry. Moreover, the majority of extant species inhabit the ancestral niche. Evidence against niche conservatism is the fact that the tribe dispersed into the tropics six separate times, which suggests that niche evolution is not rare in this lineage. However, there are two major considerations when examining these shifts. First, none of these shifts have been followed by an increase in diversification rate, which has left most of these tropical clades depauperate. Second, the ecology of some of these lineages belies their tropical distribution. For example, the Middle American clade of *Stenamma* has radiated in montane Mesoamerica, likely because of its ancestral adaptation to cool environments. Thus, while the tribe has shifted into the tropics multiple times, they have not been very successful at radiating into the niches occupied by most tropical ants, possibly due to competition.

Lastly, the diversification rate hypothesis predicts that differences in species richness are due to differences in speciation and extinction rates among clades. In the context of the standard LDG, the idea is that tropical clades will have higher speciation or lower extinction rates compared to temperate clades. However, in the case of an inverse LDG the opposite should be true. Thus, for Stenammini, we expect to see a higher diversification rate in temperate lineages. Overall this prediction is confirmed. Specifically, we found a significant increase in the diversification rate of the “Aphaenogaster” clade, a lineage with mostly temperate species. Moreover, the average speciation rate for temperate species was significantly higher than for tropical species. It should be noted, however, that with the “Aphaenogaster” clade excluded, the diversification rates among the remaining lineages, tropical or temperate, were similar.

To summarize, the predictions made by all three hypotheses are supported by our results. Stenammini has had more time to diversify in the temperate zone, it has maintained its ancestral niche as the most common niche, and it has exhibited a higher diversification rate among temperate species. Thus, all three proposed factors have contributed to the inverse diversity gradient that we see today in the Stenammini.

## ACKNOWLEDGMENTS

We would like to thank the following people for donating specimens for molecular work: Phil Ward, Marek Borowiec, Brian Fisher, Omid Paknia, Steve Shattuck, Eli Sarnat, Milan Janda, Masashi Yoshimura and Bob Johnson. We thank Jeffrey Sosa-Calvo, Theo Sumnicht, Eric Youngerman, Ana Jesovnik, and Mike Lloyd for assistance with lab work. For sequencing we thank Joe DeYoung at the UCLA Neurosciences Genomics Core, Peter Schweitzer at the Cornell Genomics Facility, and Brian Dalley at the University of Utah High Throughput Genomics Facility. Lab work for this study was conducted at the Smithsonian NMNH Laboratory of Analytical Biology (LAB) and in Lynn Boh’s molecular lab at the University of Utah. Phylogenetic analyses were performed using the Smithsonian’s High-Performance Computer Cluster (Hydra) and the CIPRES Science Gateway. Brant Faircloth provided helpful suggestions on using PHYLUCE and analyzing UCE data. We thank X anonymous reviewers for helpful suggestions to the manuscript. This work was funded by NSF grant DEB-1354739 (Project ADMAC) and the Smithsonian Competitive Grants Program for Science. MGB was partially supported by a Peter Buck Postdoctoral Fellowship at the Smithsonian.

## Supporting Information

Additional Supporting Information may be found in the online version of this article:

Appendix S1. List of valid Stenammini species and their latitudinal midpoints.

Appendix S2. Voucher information for all specimens used in the phylogeny.

Appendix S3. Sequencing and assembly statistics.

Appendix S4. Results from matrix-filtering experiments.

Appendix S5. Additional trees from RAxML, ExaBayes, and ASTRAL analyses.

Appendix S6. Complete results for divergence dating, biogeography and habitat reconstruction.

Appendix S7. Additional diversification rate results.

## DATA ACCESSIBILITY

All raw sequence data are available from the NCBI Sequence Read Archive (upon acceptance). The concatenated Filter90 alignment and corresponding ML tree are available at TreeBASE (upon acceptance).

## BIOSKETCH

**Michael Branstetter** was recently a postdoctoral scholar at the Smithsonian Institution (NMNH) and is now a postdoc at the University of Utah. He is broadly interested in the evolution of ants and other social Hymenoptera, with an emphasis on the Neotropical region. His research combines phylogenetic systematics, biogeography, and macroevolution to better understand the origin and maintenance of biodiversity.

Author contributions: All authors helped design the study and revise the manuscript. M.G.B. performed all lab work and analyses and drafted the manuscript.

## REFERENCES

Aberer A.J., Kobert K., & Stamatakis A. (2014) ExaBayes: massively parallel Bayesian tree inference for the whole-genome era. Molecular Biology and Evolution, 31, 2553–2556.

De Andrade M.L. (1995) The ant genus *Aphaenogaster* in Dominican and Mexican amber (Amber Collection Stuttgart: Hymenoptera, Formicidae. IX: Pheidolini). Stuttgarter Beiträge zur Naturkunde. Serie B (Geologie und Paläontologie), 223, 1–11.

Bayzid M.S., Mirarab S., Boussau B., & Warnow T. (2015) Weighted statistical binning: enabling statistically consistent genome-scale phylogenetic analyses. PLoS ONE, 10, e0129183.

Bininda-emonds O.R.P., Cardillo M., Jones K.E., Macphee R.D.E., Beck R.M.. D., Grenyer R., Price S.A., Vos R.A., Gittleman J.L., & Purvis A. (2007) The delayed rise of present-day mammals. Nature, 446, 507–512.

Boer P. (2013) Revision of the European ants of the Aphaenogaster testaceopilosa-group (Hymenoptera: Formicidae). Tijdschrift voor Entomologie, 156, 57–93.

Bolton B. (1982) Afrotropical species of the myrmicine ant genera Cardiocondyla, Leptothorax, Melissotarsus, Messor and Cataulacus (Formicidae). Bulletin of the Brittish Museum of Natural History (Entomology), 45, 307–370.

Bolton B. (2015) An online catalog of the ants of the world. Available at www.http://antcat.org. [Date Accessed: 2015-12-01]

Borowiec L. & Salata S. (2013) Ants of Greece – additions and corrections (Hymenoptera: Formicidae). Genus, 24, 335–401.

Borowiec L. & Salata S. (2014) Review of Mediterranean members of the Aphaenogaster cecconii group (Hymenoptera: Formicidae), with description of four new species. Zootaxa, 3861(1), 40–60.

Brady S.G., Schultz T.R., Fisher B.L., & Ward P.S. (2006) Evaluating alternative hypotheses for the early evolution and diversification of ants. Proceedings of the National Academy of Sciences, 103, 18172–18177.

Branstetter M.G. (2009) The ant genus Stenamma Westwood (Hymenoptera: Formicidae) redefined, with a description of a new genus Propodilobus. Zootaxa, 2221, 41–57.

Branstetter M.G. (2012) Origin and diversification of the cryptic ant genus Stenamma Westwood (Hymenoptera: Formicidae), inferred from multilocus molecular data, biogeography and natural history. Systematic Entomology, 37, 478–496.

Branstetter M.G. (2013) Revision of the Middle American clade of the ant genus Stenamma Westwood (Hymenoptera, Formicidae, Myrmicinae). ZooKeys, 295, 1–277.

Brown J.H. (2014) Why are there so many species in the tropics? Journal of Biogeography, 41, 8–22.

Burgess N.D., Clarke G.P., & Rodgers W.A. (1998) Coastal forests of eastern Africa: status, endemism patterns and their potential causes. Biological Journal of the Linnean Society, 64, 337–367.

Cardillo M., Orme C.D.L., & Owens I.P.F. (2005) Testing for latitudinal bias in diversification rates: An example using new world birds. Ecology, 86, 2278–2287.

Carpenter F.M. (1930) The fossil ants of North America. Bulletin of the Museum of Comparative Zoology, 70, 1–66.

Castresana J. (2000) Selection of conserved blocks from multiple alignments for their use in phylogenetic analysis. Molecular Biology and Evolution, 17, 540–552.

Chown S.L., Gaston K.J., & Williams P.H. (1998) Global patterns in species richness of pelagic seabirds: the Procellariiformes. Ecography, 21, 342–350.

Couvreur T.L.P., Chatrou L.W., Sosef M.S.M., & Richardson J.E. (2008) Molecular phylogenetics reveal multiple tertiary vicariance origins of the African rain forest trees. BMC biology, 6, 54.

Demarco B.B. & Cognato A.I. (2015) Phylogenetic analysis of Aphaenogaster supports the resurrection of Novomessor (Hymenoptera: Formicidae). Annals of the Entomological Society of America, 108, 201–210.

Dixon A.F.G., Kindlmann P., Leps J., & Holman J. (1987) Why there are so few species of aphids, especially in the tropics. The American Naturalist, 129, 580–592.

Donoghue M.J. (2008) A phylogenetic perspective on the distribution of plant diversity. Proceedings of the National Academy of Sciences, 105, 11549–11555.

Drummond A.J., Suchard M.A., Xie D., & Rambaut A. (2012) Bayesian phylogenetics with BEAUti and the BEAST 1.7. Molecular Biology and Evolution, 29, 1969–1973.

Eberle J.J. & Greenwood D.R. (2012) Life at the top of the greenhouse Eocene world: a review of the Eocene flora and vertebrate fauna from Canada’s High Arctic. Geological Society of America Bulletin, 124, 3–23.

Faircloth B.C. (2015) PHYLUCE is a software package for the analysis of conserved genomic loci. Bioinformatics, **Advance Access**, 1–3.

Faircloth B.C., Branstetter M.G., White N.D., & Brady S.G. (2015) Target enrichment of ultraconserved elements from arthropods provides a genomic perspective on relationships among Hymenoptera. Molecular Ecology Resources, 15, 489–501.

Faircloth B.C. & Glenn T.C. (2012) Not all sequence tags are created equal: designing and validating sequence identification tags robust to indels. PloS ONE, 7, e42543.

Faircloth B.C., McCormack J.E., Crawford N.G., Harvey M.G., Brumfield R.T., & Glenn T.C. (2012) Ultraconserved elements anchor thousands of genetic markers spanning multiple evolutionary timescales. Systematic Biology, 61, 1–10.

Field R., Hawkins B.A., Cornell H. V., Currie D.J., Diniz-Filho J.A.F., Guégan J.-F., Kaufman D.M., Kerr J.T., Mittelbach G.G., Oberdorff T., O’Brien E.M., & Turner J.R.G. (2009) Spatial species-richness gradients across scales: a meta-analysis. Journal of Biogeography, 36, 132–147.

Frandsen P.B., Calcott B., Mayer C., & Lanfear R. (2015) Automatic selection of partitioning schemes for phylogenetic analyses using iterative k-means clustering of site rates. BMC Evolutionary Biology, 15, 1–17.

Grabherr M.G., Haas B.J., Yassour M., Levin J.Z., Thompson D.A., Amit I., Adiconis X., Fan L., Raychowdhury R., Zeng Q., Chen Z., Mauceli E., Hacohen N., Gnirke A., Rhind N., di Palma F., Birren B.W., Nusbaum C., Lindblad-Toh K., Friedman N., & Regev A. (2011) Full-length transcriptome assembly from RNA-Seq data without a reference genome. Nature Biotechnology, 29, 644–52.

Graham A. (1993) History of the vegetation: Cretaceous (Maastichtian)-Tertiary. Flora of North America North of Mexico, Vol. 1 (ed. by F. of N.A.E. Committe), pp. 57–70. Oxford University Press, New York.

Graham A. (2011) The age and diversification of terrestrial new world ecosystems through Cretaceous and Cenozoic time. American Journal of Botany, 98, 336–351.

Guénard B., Economo E.P., Weiser M.D., Gomez K., & Narula N. (2015) antmaps.org. Available at: http://antmaps.org/about.html. [Date Accessed: 2015-12-01].

Harrington G.J., Eberle J., Le-Page B.A., Dawson M., & Hutchison J.H. (2012) Arctic plant diversity in the Early Eocene greenhouse. Proceedings of the Royal Society B, 279, 1515–1521.

Hawkins B.A. (2001) Ecology’s oldest pattern? Trends in Ecology & Evolution, 16, 470.

Hillebrand H. (2004) On the generality of the latitudinal diversity gradient. The American Naturalist, 163, 192–211.

Hölldobler B. & Wilson E.O. (1990) The Ants. Belknap Press, Cambridge, MA.

Jablonski D., Kaustuv R., & Valentine J.W. (2006) Out of the tropics: evolutionary dynamics of the latitudinal diversity gradient. Science, 314, 102–106.

Jaramillo C., Ochoa D., Contreras L., Pagani M., Romero M., Quiroz L., Rodriguez G., Rueda M.J., de la Parra F., Morón S., Green W., Bayona G., Montes C., Quintero O., Ramirez R., Mora G., Schouten S., Bermudez H., Navarrete R., Parra F., Alvarán M., Osorno J., Crowley J., Valencia V., & Vervoort J. (2010) Effects of rapid global warming at the Paleocene-Eocene boundary on Neotropical vegetation. Science, 330, 957–962.

Johnson R.A. (2001) Biogeography and community structure of North American seed-harvester ants. Annual Review of Entomology, 46, 1–29.

Kaspari M., Yuan M., & Alonso L. (2003) Spatial grain and the causes of regional diversity gradients in ants. The American Naturalist, 161, 459–477.

Katoh K., Misawa K., Kuma K., & Miyata T. (2002) MAFFT: a novel method for rapid multiple sequence alignment based on fast Fourier transform. Nucleic Acids Research, 30, 3059–66.

Kiran K., Alipanah H., & Paknia O. (2013) A new species of the ant genus Aphaenogaster Mayr (Hymenoptera: Formicidae) from Iran. Asian Myrmecology, 5, 45–51.

Krug A.Z., Jablonski D., & Valentine J.W. (2007) Contrarian clade confirms the ubiquity of spatial origination patterns in the production of latitudinal diversity gradients. Proceedings of the National Academy of Sciences, 104, 18129–18134.

Lavin M. & Luckow M. (1993) Origins and relationships of tropical North America in the context of the boreotropics hypothesis. American Journal of Botany, 80, 1–14.

Mannion P.D., Upchurch P., Benson R.B.J., & Goswami A. (2013) The latitudinal biodiversity gradient through deep time. Trends in Ecology & Evolution, 29, 42–50.

Mirarab S., Reaz R., Bayzid M.S., Zimmermann T., Swenson M.S., & Warnow T. (2014) ASTRAL: genome-scale coalescent-based species tree estimation. Bioinformatics, 30, i541–i548.

Moreau C.S. & Bell C.D. (2013) Testing the museum versus cradle tropical biological diversity hypothesis: phylogeny, diversification, and ancestral biogeographic range evolution of the ants. Evolution, 67, 2240–2257.

Olson D.M., Dinerstein E., Wikramanayake E.D., Burgess N.D., Powell G.V.N., Underwood E.C., D’amico J.A., Itoua I., Strand H.E., Morrison J.C., Loucks C.J., Allnutt T.F., Ricketts T.H., Kura Y., Lamoreux J.F., Wettengel W.W., Hedao P., & Kassem K.R. (2001) Terrestrial ecoregions of the world: a new map of life on Earth. BioScience, 51, 933–938.

Owen D.F. & Owen J. (1974) Species diversity in temperate and tropical Ichneumonidae. Nature, 249, 583–584.

Paradis E., Claude J., & Strimmer K. (2004) APE: Analyses of Phylogenetics and Evolution in R language. Bioinformatics, 20, 289–290.

Pellmyr O. & Leebens-Mack J. (1999) Forty million years of mutualism: evidence for Eocene origin of the yucca-yucca moth association. Proceedings of the National Academy of Sciences, 96, 9178–9183.

Pianka E.R. (1966) Latitudinal gradients in species diversity: a review of concepts. The American Naturalist, 100, 33–46.

Plummer M., Best N., Cowles K., & Vines K. (2006) CODA: Convergence Diagnosis and Output Analysis for MCMC. R News, 6, 7–11.

Pyron R.A. & Burbrink F.T. (2009) Can the tropical conservatism hypothesis explain temperate species richness patterns? An inverse latitudinal biodiversity gradient in the New World snake tribe Lampropeltini. Global Ecology and Biogeography, 18, 406–415.

Rabosky D.L. (2014) Automatic detection of key innovations, rate shifts, and diversity-dependence on phylogenetic trees. PloS one, 9, e89543.

Rabosky D.L., Donnellan S.C., Grundler M., & Lovette I.J. (2014a) Analysis and visualization of complex macroevolutionary dynamics: an example from Australian scincid lizards. Systematic Biology, 63, 610–627.

Rabosky D.L., Grundler M., Anderson C., Title P., Shi J.J., Brown J.W., Huang H., & Larson J.G. (2014b) BAMMtools: an R package for the analysis of evolutionary dynamics on phylogenetic trees. Methods in Ecology and Evolution, 5, 701–707.

Rabosky D.L., Santini F., Eastman J., Smith S.A., Sidlauskas B., Chang J., & Alfaro M.E. (2013) Rates of speciation and morphological evolution are correlated across the largest vertebrate radiation. Nature Communications, 4, 1958.

Rambaut A., Suchard M.A., Xie D., & Drummond A.J. (2014) Tracer v1.6, Available from http://beast.bio.ed.ac.uk/Tracer.

Ree R.H. & Smith S.A. (2008) Maximum likelihood inference of geographic range evolution by dispersal, local extinction, and cladogenesis. Systematic Biology, 57, 4–14.

Rivadeneira M.M., Alballay A.H., Villafaña J.A., Raimondi P.T., Blanchette C.A., & Fenberg P.B. (2015) Geographic patterns of diversification and the latitudinal gradient of richness of rocky intertidal gastropods: the “into the tropical museum” hypothesis. Global Ecology and Biogeography, 24, 1149–1158.

Rivadeneira M.M., Thiel M., González E.R., & Haye P.A. (2011) An inverse latitudinal gradient of diversity of peracarid crustaceans along the Pacific Coast of South America: out of the deep south. Global Ecology and Biogeography, 20, 437–448.

Rohde K. (1992) Latitudinal gradients in species-diversity: the search for the primary cause. Oikos, 65, 514–527.

Salata S. & Borowiec L. (2015) A taxonomic revision of the genus Oxyopomyrmex André, 1881 (Hymenoptera: Formicidae). 4025, 1–66.

Schluter D. (1993) Species diversity: historical and geographical perspectives. University of Chicago Press, Chicago.

Seo T.K. (2008) Calculating bootstrap probabilities of phylogeny using multilocus sequence data. Molecular Biology and Evolution, 25, 960–971.

Shi J.J. & Rabosky D.L. (2015) Speciation dynamics during the global radiation of extant bats. Evolution, 69, 1528–1545.

Smith S.A., Stephens P.R., & Wiens J.J. (2005) Replicate patterns of species richness, historical biogeography, and phylogeny in Holarctic treefrogs. Evolution, 59, 2433–2450.

Stadler T. (2011) Mammalian phylogeny reveals recent diversification rate shifts. Proceedings of the National Academy of Sciences, 108, 6187–6192.

Stamatakis A. (2014) RAxML version 8: A tool for phylogenetic analysis and post-analysis of large phylogenies. Bioinformatics, 30, 1312–1313.

Stephens P.R. & Wiens J.J. (2003) Explaining species richness from continents to communities: the time-for-speciation effect in emydid turtles. The American Naturalist, 161, 112–128.

Stevens R.D. (2006) Historical processes enhance patterns of diversity along latitudinal gradients. Proceedings of the Royal Society B, 273, 2283–2289.

Talavera G. & Castresana J. (2007) Improvement of phylogenies after removing divergent and ambiguously aligned blocks from protein sequence alignments. Systematic Biology, 56, 564–577.

R Core Team (2015) R: A language and environment for statistical computing. R Foundation for Statistical Computing, Vienna, Austria.

Terayama M. (2009) A synopsis of the family Formicidae of Taiwan (Insecta, Hymenoptera). Research Bulletin of Kanto Gakuen University. Liberal Arts., 17, 81–266.

Tiffney, Bruce H. (1985) The Eocene North Atlantic land bridge: its importance in Tertiary and modern phytogeography of the Northern Hemisphere. Journal of the Arnold Arboretum, 66, 243–273.

Ward P.S., Brady S.G., Fisher B.L., & Schultz T.R. (2015) The evolution of myrmicine ants: phylogeny and biogeography of a hyperdiverse ant clade (Hymenoptera: Formicidae). Systematic Entomology, 40, 61–81.

Weir J.T. & Schluter D. (2007) The latitudinal gradient in recent speciation and extinction rates of birds and mammals. Science, 315, 1574–1576.

Wheeler W.M. (1915) The ants of the Baltic Amber. Schriften der Physikalisch-Ökonomischen Gesellschaft zu Königsberg, 55, 1–142.

Whitford W.G., Depree E., & Johnson P. (1980) Foraging ecology of two Chihuahuan desert ant species: Novomessor cockerelli and Novomessor albisetosus. Insectes Sociaux, 27, 148–156.

Wiens J.J. (2007) Global patterns of diversification and species richness in amphibians. The American Naturalist, 170, S86–106.

Wiens J.J. & Donoghue M.J. (2004) Historical biogeography, ecology and species richness. Trends in Ecology and Evolution, 19, 639–644.

Wiens J.J. & Graham C.H. (2005) Niche conservatism: integrating evolution, ecology, and conservation biology. Annual Review of Ecology, Evolution, and Systematics, 36, 519–539.

Willig M.R., Kaufman D.M., & Stevens R.D. (2003) Latitudinal gradients of biodiversity: pattern, process, scale, and synthesis. Annual Review of Ecology, Evolution, and Systematics, 34, 273–309.

Wing S.L. (1987) Eocene and Oligocene floras and vegetation of the rocky mountains. Annals of the Missouri Botanical Garden, 74, 748–784.

Wolfe J.A. (1975) Some aspects of plant geography of the northern hemisphere during the Late Cretaceous and Tertiary. Annals of the Missouri Botanical Gardens, 62, 264–279.

Yoder A.D. & Nowak M.D. (2006) Has vicariance or dispersal been the predominant biogeographic force in Madagascar? Only time will tell. Annual Review of Ecology, Evolution, and Systematics, 37, 405–431.

Yu Y., Harris A.J., Blair C., & He X. (2015) RASP (Reconstruct Ancestral State in Phylogenies): a tool for historical biogeography. Molecular Phylogenetics and Evolution, 87, 46–49.

Zachos J., Pagani M., Sloan L., Thomas E., & Billups K. (2001) Trends, rhythms, and aberrations in global climate 65 Ma to present. Science, 292, 686–693.

